# Global remapping emerges as the mechanism for renewal of context-dependent behavior in a reinforcement learning model

**DOI:** 10.1101/2023.10.27.564433

**Authors:** David Kappel, Sen Cheng

## Abstract

The hippocampal formation exhibits complex and context-dependent activity patterns and dynamics, e.g., place cell activity during spatial navigation in rodents or remapping of place fields when the animal switches between contexts. Furthermore, rodents show context-dependent renewal of extinguished behavior. However, the link between context-dependent neural codes and context-dependent renewal is not fully understood. We use a reinforcement learning agent based on deep neural networks to study the learning dynamics that occur during spatial learning and context switching in a simulated ABA extinction and renewal paradigm in a 3D virtual environment. Despite its simplicity, the network exhibits a number of features typically found in the CA1 and CA3 regions of the hippocampus. A significant proportion of neurons in deeper layers of the network are tuned to a specific spatial position of the agent in the environment - similar to place cells in the hippocampus. These spatial representations exhibit global remapping when the agent is exposed to a new context. The spatial maps are restored when the agent returns to the previous context, accompanied by renewal of the conditioned behavior. Remapping is facilitated by memory replay of experiences during training. These complex spatial representations and dynamics occur spontaneously in the hidden layer of a deep network during learning. Our results show that integrated codes that jointly represent spatial and task-relevant contextual variables are the mechanism underlying renewal in a simulated DQN agent.

## 1 Introduction

Classical Pavlovian conditioning has taught us that animals can learn to emit a conditioned response (CR) to a conditioned stimulus (CS) when the CS is repeatedly paired with a physiologically relevant, unconditioned stimulus (US), which usually involves reinforcement or punishment (Pavlov, 1927). When the CS is subsequently presented repeatedly without the US, the association between CR and CS weakens, a phenomenon referred to as extinction learning (Auchter et al., 2017). However, this effect has been found to be complex and highly context-dependent, which can lead to renewal of learned behavior under certain circumstances (Dunsmoor et al., 2015).

This effect is most evident in the ABA renewal paradigm (Bouton, 2004; Corcoran, 2004; Ji and Maren, 2008; Fujiwara et al., 2012; Zelikowsky et al., 2013). In this experimental design, a subject typically acquires a CR to a CS when it is paired with an US in a particular context A, e.g., defined by a spatial enclosure, an odor or light stimulus, and then the response is extinguished in a different context B in the absence of the US. When the subject returns to context A (without the US), the CR is restored. The ABA renewal phenomenon suggests that extinction learning does not completely erase or overwrite the previously learned association, but rather forms a new contextual association (Dunsmoor et al., 2015).

The hippocampal formation plays an important role in extinction learning (Jones et al., 2016; Bernier et al., 2017; Hainmueller and Bartos, 2020), and it was found that extinction learning depends on learning mechanisms in the hippocampus (Peters et al., 2010; Soliman et al., 2010; Rosas-Vidal et al., 2014; Wang et al., 2018; Bouton et al., 2021). Furthermore, the hippocampal formation shows context-dependent activity patterns and complex dynamics, e.g. place cell activity during spatial navigation in rodents or place field remapping when animals are moved between different environments (Grieves and Jeffery, 2017; Latuske et al., 2018) (Fig. 1A). The latter phenomenon is referred to as global remapping and means that some place cells are active only on one environment, and not the other, and some place cells have place fields in different relative locations in the two environments (Leutgeb et al., 2005). Hippocampal place field maps also encode abstract task-relevant variables (Knudsen and Wallis, 2021). Some authors have therefore proposed that remapping is the physiological basis for task-relevant context variables (Kubie et al., 2020; Plitt and Giocomo, 2021; Sanders et al., 2020), i.e., that place field maps jointly encode place and context in an integrated code (Fig. 1C).

**Figure 1:**
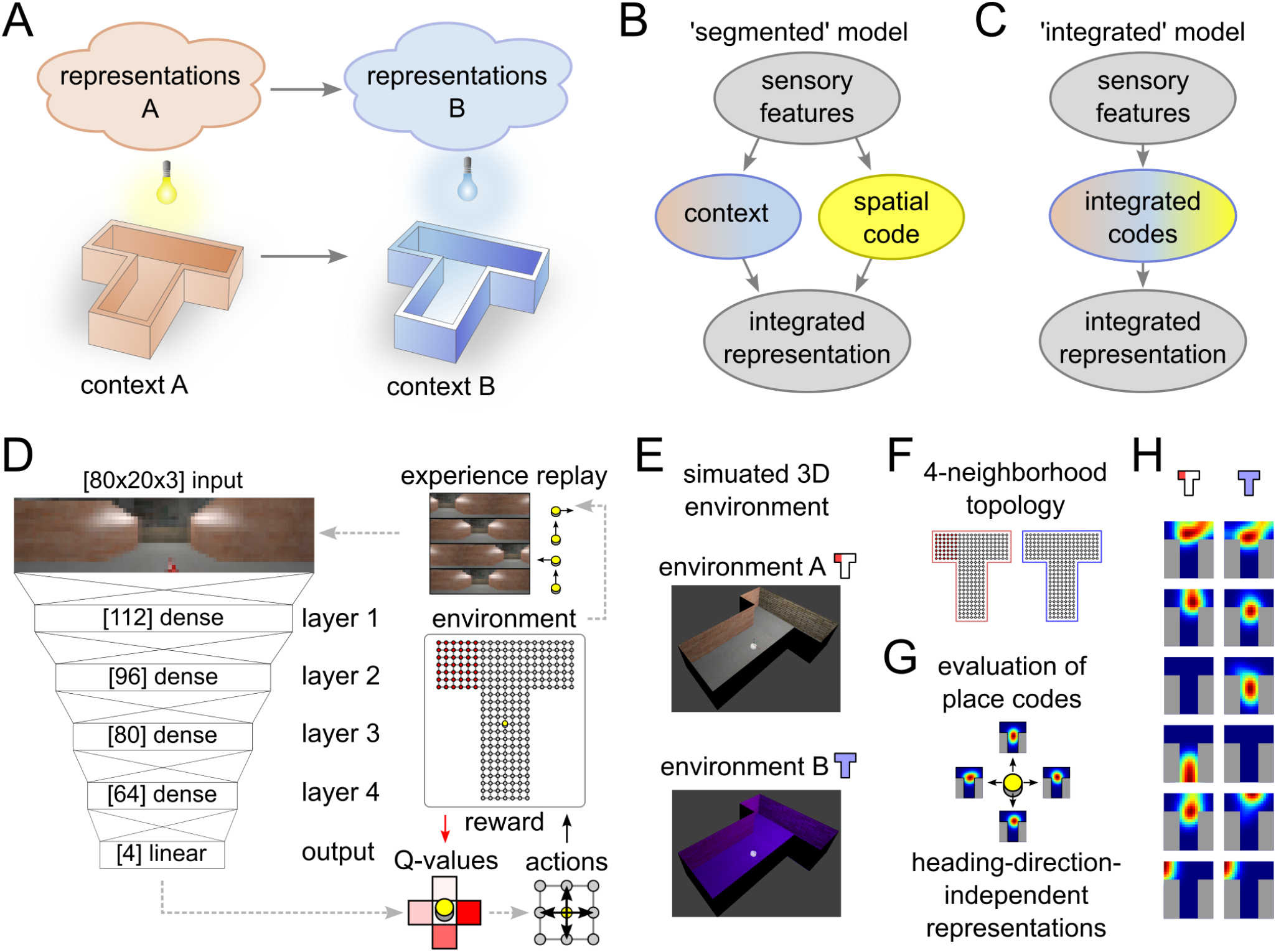
Experimental setup and data analysis. **A:** Illustration of remapping paradigm. Moving from environment A to B results in the expression of a different spatial representation. **B,C:** Two alternative models that can explain place field remapping, the ‘segmented’ model (B) separates codes for space and context, while the ‘integrated’ model (C) combines these codes. **D,E:** Illustration of the reinforcement learning agent (D) that interacts with a simulated environment (E). **F-H:** Place field analysis performed on the neural activations. 4-neighborhood topology in the T-maze (F) is used to record neural activity maps for each heading direction (G). Activity of representative example neurons that showed place-cell-like behavior in one or both environments.

Modern machine learning techniques, are now able to solve behavioral tasks that are comparable in complexity to real-world behavior and to reach human performance in some specific domains (Ashraf et al., 2021). These techniques have been used to model hippocampus functionality on an abstract level (Diekmann and Cheng, 2023) and to explain the emergence of complex task-relevant spatial codes (Vijayabaskaran and Cheng, 2022). Furthermore, in Walther et al. (2021), a simplified abstract ABA extinction learning paradigm was tested in a simulated agent. It was found that the prominent signs of extinction and renewal could be reproduced in the model, but the underlying representations of space and context that emerged in the circuit were not studied systematically.

A number of models for remapping have been proposed. An early model that focused on place cell remapping described the phenomenon as attractor dynamics between spatially tuned neurons (McNaughton et al., 1996; Samsonovich and McNaughton, 1997), but assumed the existence of special signals to represent cues and contexts, and did not learn from realistic visual input. Other earlier models proposed to view the hippocampus as a hierarchical generative model (Stoianov et al., 2021; Taniguchi et al., 2022; Penny et al., 2013). Following these models, context can be recovered from network activity through statistical inference (Fuhs and Touretzky, 2007). A related idea also underlies an earlier statistical model of memory stability (Gershman et al., 2017). Based on these results, a general contextual inference theory of sensory-motor learning has recently been proposed (Heald et al., 2021). Redish et al. (2007) proposed a simplified computational model for extinction and renewal based on the explicit representation of contextual cues.

In this study, we develop a simulation environment that uses naturalistic inputs with implicit contextual information to study the emergence of representations in an ABA extinction and renewal paradigm. We show the emergence of context-dependent spatial codes that closely resemble place cells, as well as a context-dependent remapping of these codes in the simulated agent. We compare two alternative hypotheses about the encoding of context in complex behaving agents: A segmented code would suggest that separate populations of neurons encode context and spatial location (Fig. 1B), whereas an integrated code would suggest mixed encoding of space and context (Fig. 1C). We show the simultaneous emergence of spatial representations and task-relevant context variables throughout all layers of the reinforcement learning agent. These emerging representations in our model are more compatible with the integrated than with the segmented model, which is compatible with neural codes in the hippocampus. Furthermore, we show that the formation of independent spatial maps depends on the salience of the context stimulus and investigate the role of experience replay and find a strong dependence between renewal and replay.

## 2 Methods

### 2.1 Simulated behavioral experiment

To test the behavior and spatial representations that emerge in the simulated agent during an ABA renewal paradigm, we designed a closed-loop simulation of a spatial task inspired by forced choice T-maze experiments (see Fig. 1D-F). In every trial, the agent was located at the bottom center of the base of the T-maze and could move freely between grid points (238 in total) by choosing to move in one of four cardinal directions (North, East, South, West). Actions that would result in a movement outside the grid world consumed one time step but otherwise had no effect. Heading directions of the agent were rotated in alignment of the actions in every time step. When a goal location (red) was visited, the agent received a reward (+20) and the trial was terminated. In every other time step, a negative reward of -1 was given to encourage the agent to find the goal as fast as possible. Unsuccessful trials were terminated after 400 time steps.

All simulations were based on the CoBeL-RL simulation framework (Diekmann et al., 2023). Visual feedback was provided to the artificial agent in the form of naturalistic images (Fig. 1E) generated using the Blender 3D computer graphics software (version 2.79). The maze had a length of 2.8 meters and walls were structured with photo-realistic wall textures. A 360°view image of a typical lab environment was projected onto a cylindrical sky box around the maze to provide distal spatial cues. A point light source with adjustable color was placed above the center of the T-maze to control the context (blue and white light condition). To generate the rendered input images, a camera object was placed at the agent’s location and heading direction. Gaussian jitter with standard deviations 1 cm and 10°were added to the camera position and orientation, respectively. An image was then captured, scaled and cropped to 80 *×* 20 pixels, corresponding to an effective horizontal angle of view of around 240°.

### 2.2 Deep Q-learning Network

To model spatial learning, we adopted reinforcement learning and, since the goal of this study was to analyze emergent spatial representations, we employed a Deep-Q Learning Network (DQN) agent. The DQN agent combines deep neural networks (DNNs) and Q-learning, i.e. the action-value function *Q*(*s, a*) is represented and approximated by a DNN (Ashraf et al., 2021). The update scheme of *Q* realizes one iteration of the Bellman equation, i.e. for the current state *s*, action *a*, reward *r*, and next state *s*^*′*^

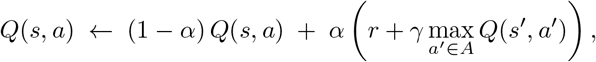

where *α* is a learning rate, *γ* is the reward discount factor and *A* is the state space of the actions. The Adam optimizer was used during training to control the learning dynamics, with a learning rate parameter of 10^*−*4^.

Inside the DQN agent, a DNN was used to infer Q-values in response to complex visual stimuli. The DNN architecture in our agent consisted of a feed-forward densely-connected artificial neural network with 4 hidden layers (Fig. 1D). Hidden layer sizes from input to output were 112, 96, 80 and 64. Batch normalization followed by rectified linear activation functions were used to create layer outputs. The learning rule and network architecture allowed us to study the spatial representations that emerge during learning the the ABA renewal paradigm.

Experience replay was used during training by sampling experiences consisting of (*s, a, r, s*^*′*^) tuples randomly from a replay buffer, if not stated otherwise. In the condition without experience replay (Fig. 5), experiences were randomly sampled from the last 5 trials for every training epoch.

The *ε*-greedy strategy was used to drive exploration during training, i.e., in every time step, the action corresponding to the highest Q-value was chosen with probability 1*− ϵ* and a random action with probability *ϵ. ϵ* was chosen to be 0.3 in all experiments.

### 2.3 Behavioral analysis

Behavioral trajectories (sequence of grid points visited in the maze) were recorded in all experiments. Figure 2A shows the superposition of 50 successive movement trajectories. Every trial was classified based on these trajectories as conditioned response (CR) trial or non-CR trials. Grid point location trajectories of the agent were captured throughout the whole experiment. For CR analysis, agent locations were tracked over single trials. Trials were counted as CR trials if the agent entered the left arm of the maze within the first 50 time steps. Learning dynamics were visualized using cumulative response curves (CRC) (Donoso et al., 2021). CRC plots in Fig. 2 show averages over 20 independent experimental runs.

**Figure 2:**
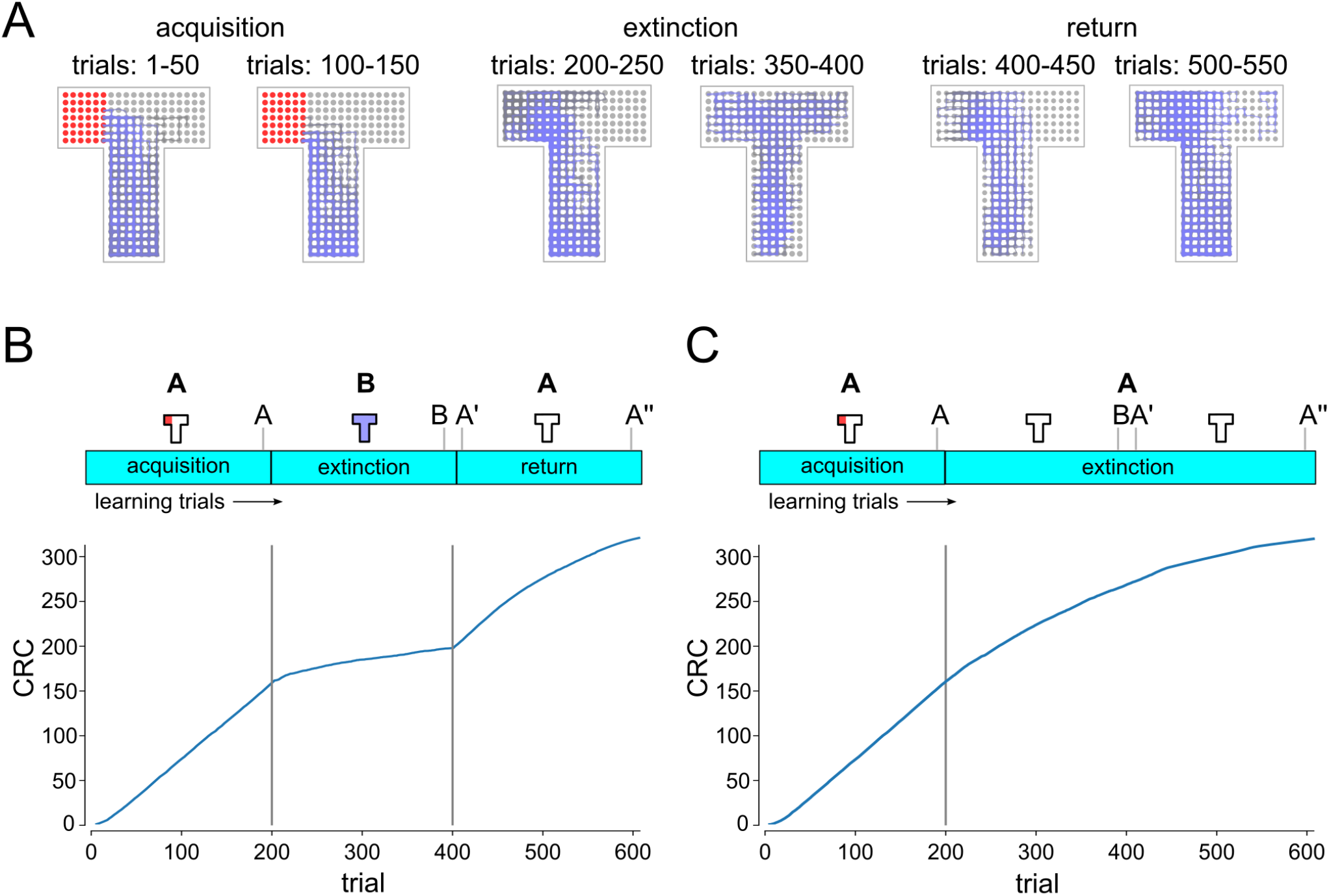
Analysis of the behavior of the simulated DQN agent. **A:** Superposition of movement trajectories at different stages of the experiment. Colors indicate relative trial index (gray: earlier, blue: later). Reward locations indicated in red. Extinction and renewal in simulated agent. Cumulative response curves for **B:** ABA renewal and **C:** extinction experiment. Progression of extinction and renewal tasks and population vector recording time points (A,B,A’,A”) are indicated on top. Renewal of conditioned behavior is clearly visible after return to A in ABA task (B) but not in AA task (C).

### 2.4 Place field analysis

To analyze emergent spatial representations (see Fig. 1F-H for an illustration) we adopted the method from Vijayabaskaran and Cheng (2022). First, activity vectors were recorded from all neurons of hidden layers in the network by placing the agent at all possible grid point locations and heading directions. Cells that produced zero output under all condition were classified as silent cells. Cells that produced non-zero activity at some locations for at least one, but not all, heading directions were classified as partially active neurons.

Preferred firing centers were further analyzed for the remaining neurons. Activity vectors were smoothed with a Gaussian smoothing filter before a cluster analysis was performed for each heading direction. Neurons with activity that revealed more than 2 clusters or neurons that were active at more than 50% of the grid point locations were excluded from the analysis. Heading-direction specific tuning centers were then computed as the weighted average of grid point locations in the dominating activity cluster. Cells for which the heading-direction specific tuning center exceeded the distance of 4 times the distance between grid points, were excluded. All neurons that were excluded in the above tests were classified as heading-direction modulated cells. The remaining neurons were classified as place-cell-like.

For the remapping analysis in Fig. 3, only neurons that were classified as place-cell-like in at least one of the experimental conditions were examined. Population vectors (PVs) were generated by concatenating the activity vectors of these neurons, merging the per-grid-point activities for the four heading directions into one. PVs were pooled together from 20 independent runs. Correlation coefficients were then computed on these PV amplitudes.

**Figure 3:**
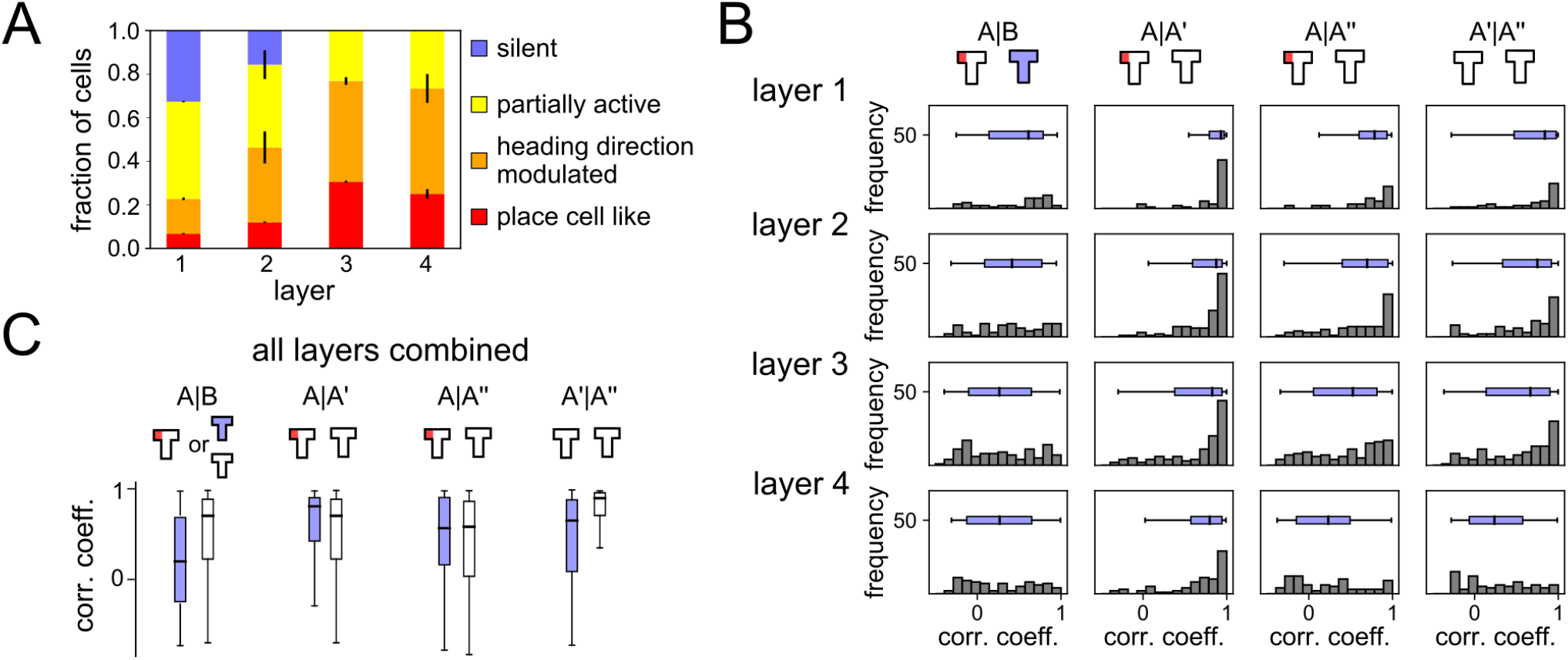
Emergent spatial representations and remapping. **A:** Distribution of different cell types that emerged the different network layers during learning in context A. Deeper layers developed spatial coding more frequently. **B:** Histograms of correlation coefficients between population vectors recorded at different phases of the ABA renewal paradigm. All layers show signs of global remapping, i.e., spatial correlations are low between contexts A and B, but high between two different exposures to the same context A. **C:** Comparison of PV correlation coefficients for the ABA renewal paradigm (blue) and AA extinction (white). Data from all network layers are combined. Without the context change spatial representations remain stable throughout extinction.

## 3 Results

### 3.1 Extinction and renewal dynamics

Walther et al. (2021) demonstrated that extinction and renewal can be reproduced in an *in-silico* model, using a simplified abstract ABA extinction learning paradigm. We first studied whether extinction and renewal behavior could be reproduced in a more complex simulated agent that interacts with a 3D environment. To do so, we designed a T-maze paradigm with a fixed reward location as the basis for the ABA renewal experiment (Fig. 2). The agent is first placed in environment A (T-maze with reward zone in the left arm, white light) for 200 trials, before the behavior is extinguished in environment B in context B (T-maze without reward, blue light). We recorded behavioral trajectories and examined signs of extinction and renewal for the simulated agent in this ABA renewal paradigm.

Extinction and renewal emerged in the simulated agent when exposed to the ABA task (Fig. 2A,B). Superposition of movement trajectories at different stages of the experiment shows that the agent quickly adopts the conditioned response (CR), i.e., prefers the left arm of the maze, entering the reward zone typically within the first 50 time steps of the trial (Fig. 2A). After 200 trials, the agent was moved to context B (blue light) and reward was suspended. CR preference briefly continues after switch to B but quickly washes out to baseline showing no preference between left and right arm (see Fig. 2B). After returning to context A at trial 400 the CR is renewed followed by a slow gradual washout of CR. As in experiments, renewal requires the switch of context, i.e., if acquisition, extinction and test occur in the same context, there is no renewal (Fig. 2C).

### 3.2 Emergent remapping of spatial representations

We next analyzed the emerging neural codes in the DQN agent after it has learned this task. The neural code analysis was performed at the end of every experimental phase. Based on these recordings and using a method adapted from Vijayabaskaran and Cheng (2022) (see Sec. 2.4), neurons of the DQN agent were classified into one of the four classes: silent, partially active, heading direction modulated, and place cell like. Figure 1H shows example place fields extracted from the DQN. While place-cell-like responses could be found in all layers and experimental phases (Fig. 3A), deeper layers had a larger number of spatially tuned neurons, e.g. in layer 3 30.62 *±* 0.9 % of the units were classified as place cell like, 46.24 *±* 3.4 % as heading direction modulated, and 23.12 *±* 3.6 % as partially active. A significant number of silent cells were found in layer 1 (32.58 *±* 0.6 %) and layer 2 (15.63 *±* 13.2 %) and these layers were overall less tuned to spatial features. No cells were found that responded exclusively to the context, without being also tuned to spatial features or heading direction.

A new spatial code emerged when the agent was moved to context B as indicated by histograms of correlation coefficients for population vectors (PVs) from different experimental phases and network layers (Fig. 3B). PVs in contexts A and B were only weakly correlated (histograms are flat or skewed towards 0), whereas correlations between PVs in context A at different time points (A, A’, A”) showed strong correlations with each other. Overall deeper layers showed slightly stronger signs of remapping.

Figure 3C shows a comparison of correlation coefficients between the ABA and AA experiment. Remapping occurred in the ABA condition, whereas correlation coefficients were better explained by temporal proximity in AA condition, suggesting a slow drift of neural representations rather than remapping. Our results suggest that global remapping in the spatial representations is a key mechanism underlying renewal.

### 3.3 The role of context salience in remapping and renewal

Our results in the preceding section suggest that remapping of task-relevant representations in the network is driving renewal in the test phase. To further investigate this dependency we adapted the experiment to include a variable strength of context salience. The difference between the A and B contexts is determined by a change of illumination in the T-maze. To test the sensitivity of the emergent spatial representations to changes in the differentiability between A and B context, we repeated the experiment with different lighting conditions that blend between the A (white light) and B (blue light) context, parametrized by a new task variable (saliency of context, *s*). While context A remains fixed (white light), *s* = 0 indicates that both contexts are identical (white light in B), *s* = 1 corresponds to pure blue light in B, and values between 0 and 1 denote a mix of blue and withe light in context B. Clearly, *s* = 0 corresponds to extinction without context switch (Fig. 2C).

We repeated the analysis of remapping in neural codes for experiments with different values of *s* (0*≤ s≤* 1) and compared mean correlation coefficients between PVs from experimental phases A, B and A’ (Fig. 4A). As expected correlations of PVs are identical between A-B and A-A’ for *s* = 0 (A, A’ and B conditions are indistinguishable) and maximally different for *s* = 1, suggesting high levels of remapping between context A and B. Interestingly, we found that A-A’ correlation coefficients showed a U-shaped profile. For *s* = 0, the correlations are high because the contexts A and B are identical and so are their representations. In that situation, extinction learning overwrites the acquired behavior so that there is no renewal. If context salience is low to intermediate (0 *≤ s ≤* 0.4), training in B causes interference with the previously learned spatial representations of A. For *s >* 0.4, the two context are less similar and the interference between the spatial representations ameliorates. Despite the interference between spatial representations at low to intermediate context salience, renewal of CR could be observed already at quite low levels of context salience (Fig. 4B). In Fig. 4C we show the correlation coefficient between A’ and B, which is a measure of interference between the neural patterns needed to learn the behavior. As can be seen interference also declines much slower than the rise in CR trials in Fig. 4B. These findings suggest that at low context salience even though spatial representations are distinct enough to support different associations, their difference are not well captured by the analyzing the correlations between the PVs. For higher context salience (*s >* 0.4), the agent learns distinct representations of A and B, leading to much higher A-A’ than A-B correlations, i.e., global remapping, and renewal.

**Figure 4:**
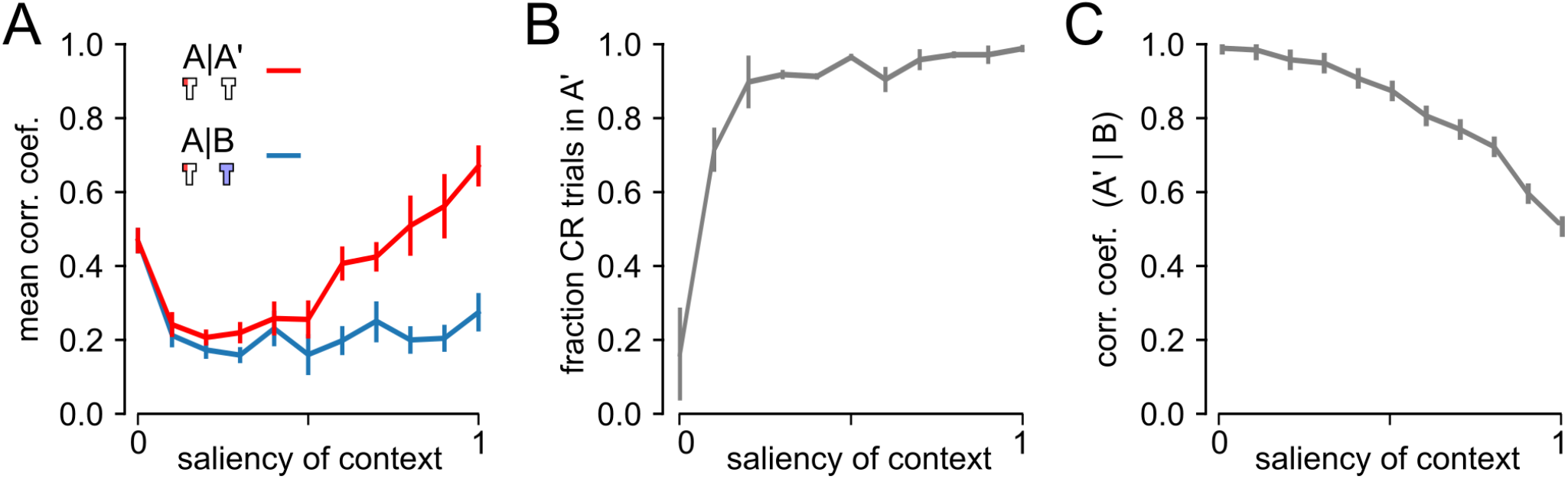
Impact of context salience on remapping and behavior. **A:** Correlation coefficient between population vectors (PVs) taken at different phases of the experiment as a function of the context salience. A vs. A’ PVs show a U-shaped profile suggesting interference between PVs for low and intermediate context salience. **B:** Fraction of CR trials in the A’ phase (error bars show STD). **C:** Correlation coefficient between A’ and B, to measure the interference between emerging neural patterns as a function of context salience.

### 3.4 Role of experience replay in remapping

We next wondered what caused remapping in the DQN agent’s neural representations. A simple hypothesis to explain the observed remapping behavior is that experience replay, that spans multiple experimental phases, rescues neural representations from forgetting and thus facilitates the emergence of two parallel spatial maps. This hypothesis would suggest that remapping would be abolished if replay were absent during training.

To test this prediction we repeated the ABA renewal experiment with maximum context salience (*s* = 1), using a DQN agent without experience replay. Emergence of place cells did not depend on replay as spatially tuned neurons also emerged in agents trained without experience replay (Fig. 5A). As expected, PV correlations are overall decreased in all experimental phases, suggesting that experience replay has a stabilizing effect on neural representations (Fig. 5B). A closer inspection of the relative correlation coefficients reveals that they are higher for A-A’ than for A-B, despite the fact that the latter are more closely spaced in time, indicating that global remapping occurs even in the no-replay condition. However, since the representations of A and B are less distinct from one another than in the simulations with replay, we conclude that replay helps in the formation of distinct representations during extinction learning in context B.

**Figure 5:**
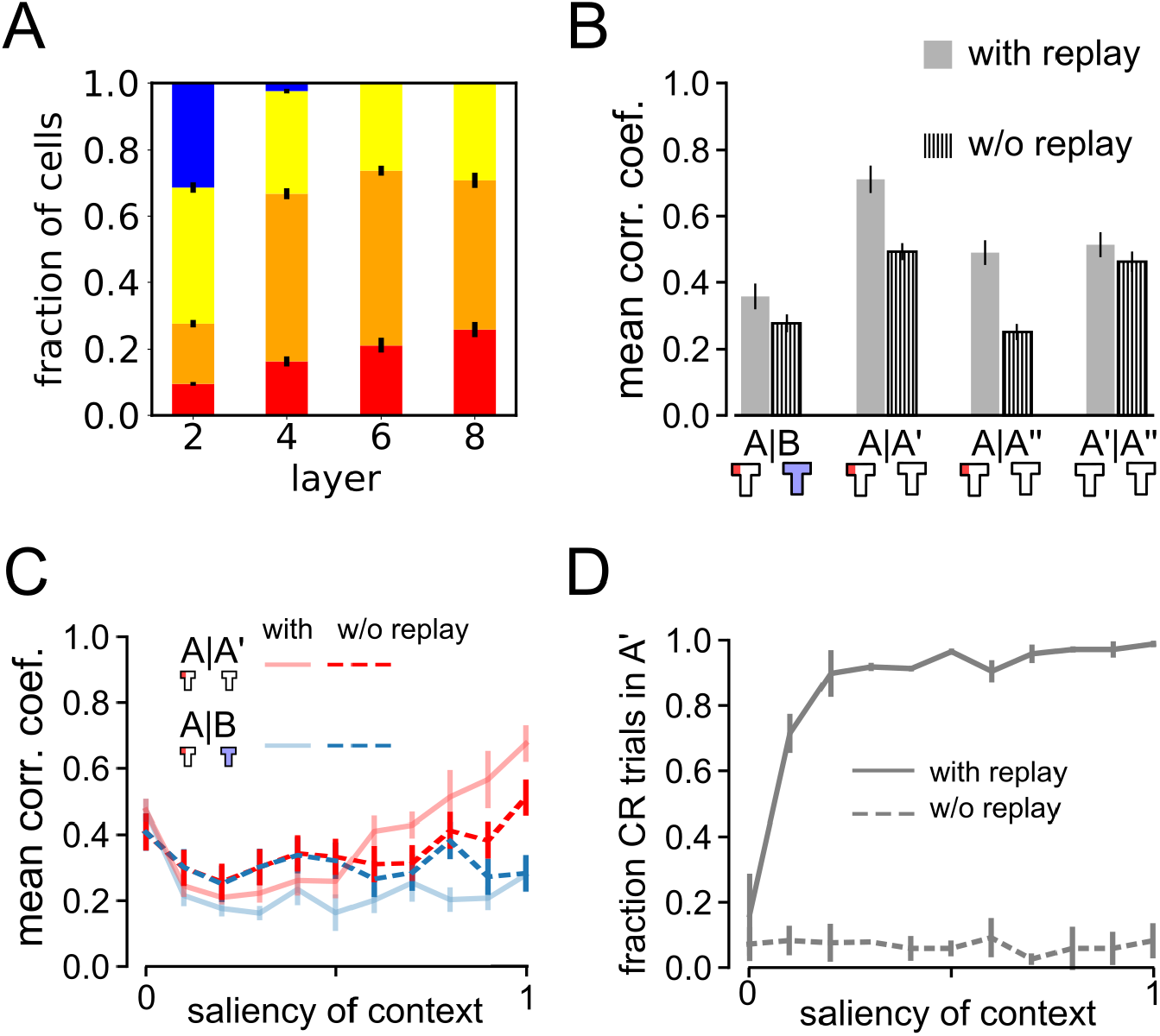
Impact of experience replay on remapping and behavior. **A:** Relative fraction of cell types as in Fig. 3A but for an agent trained without experience replay. **B:** Remapping analysis similar to Fig. 3C showing less clear remapping when replay is absent. **C,D:** Analysis of impact of context salience on remapping and behavior as in Fig. 4 but for an agent trained without experience replay. Data without experience replay displayed with dashed lines. Solid lines show data with replay as in Fig. 4 for comparison.

This conclusion is supported by another observation when comparing the representations of A, A’, and A” (Fig. 5B). There are roughly the same number of learning trials between A-A’ and A’-A”, but between A and A’ extinction occurs in context B whereas all trials between A’ and A” occur in context A. Interestingly, without replay the A-A’ correlations are similar to A’-A” correlations, suggesting that the context of the intervening trials has no influence on the stability of the representation of context A. However, if there is replay, A-A’ correlations are higher than A’-A”, where the latter is similar to the no-replay condition. This might seem counterintuitive since between A and A’ extinction occur in a different context. To explain this observation, we note that the trials in the third experimental phase are not reinforced, so extinction learning should occur, too, but in context A. It is reasonable to assume that extinction learning leads to changes in neural representations to drive a different behavior. Our results suggest that if extinction learning occurs in a different context from acquisition, replay strengthens the representation of the acquisition context.

Finally, we studied the dependence between remapping and renewal in the no-replay condition by varying the context salience *s* (Fig. 5C,D). Agents trained with experience replay (solid lines) are compared to those without replay (dashed lines). Without replay remapping is apparent only for the highest context salience and not for lower values. On the other hand, no signs of renewal were found in the agents’ behavior (Fig. 5D). In summary, our modeling results results suggest a crucial role of experience replay in renewal and an important role in remapping.

## 4 Discussion

In this study, we have shown that DQN agents trained for spatial tasks learn complex representations that encode both spatial position and contextual variables about the environment. We used an ABA renewal paradigm where a CR is extinguished in the B context. We reproduced marked signs of renewal after the agent is returned to context A across a wide range of experimental settings. We showed that complex context-dependent spatial representations and dynamics arise spontaneously in the hidden layer activity of a deep network when a spatial navigation task is learned. We found that the distinctness of contextual cues plays an important role in global remapping of spatial representations and that the occurence of global remapping strongly correlates with renewal behavior. Finally, we found that experience replay is critical to stabilize acquired behavior during the extinction phase. Disabling experience replay also prevented renewal and reduced the remapping effect.

These results demonstrate that deep-learning agents can reproduce a set of complex learning dynamics similar to those found in behaving animals. These results may therefore provide new insights to better understand how spatial representations are formed in the brain to support goal-directed behavior.

### 4.1 Plausibility of our modeling assumptions

Reinforcement learning provides a robust simulation framework for understanding the principles underlying navigation. As in biological systems an RL agent has to cope with a number of complex problems such as interpreting sensory stimuli, extracting task-relevant features form these stimuli, or dealing with the exploration-exploitation trade-off. However, since reinforcement learning is driven by the goal of reward acquisition, it neglects other intrinsic learning mechanisms, such as curiosity or fatigue.

Furthermore, the combination of deep neural networks and reinforcement learning, which forms the basis of a DQN agent, makes it possible to process naturalistic inputs. However, the DQN setup models neural activity only in an abstract way by generating analog neural outputs at each time step. These outputs model the activations of biological neurons at a coarse temporal resolution, roughly corresponding to neuronal firing rates or calcium transients. It will be interesting to study whether similar learning dynamics can be observed in more detailed models of biological neurons, such as Frémaux et al. (2013).

To maintain full control over the experimental setup we studied here a simple feed-forward DQN agent. For example, we focused on a relatively abstract interaction with the environment where only 4 different actions could be generated in every time step. By augmenting the model with more behavioral details, a number of additional experimental findings could be tested in future work. For example, experimental evidence indicates that hippocampal cognitive maps encode abstract task-relevant dimensions that encode value (Knudsen and Wallis, 2021), suggesting a complex mixed encoding of spatial representations, context, and task-relevant variables.

As for the behavior, we focused on a T-maze paradigm, where the agent learns to navigate to a fixed local target to retrieve a reward. This experimental paradigm fits well with the reinforcement learning setup, and the DQN agent was able to quickly acquire this behavior. However, experimental studies have also explored other conditions for reliably inducing global or partial remapping, such as different maze topologies (Leutgeb et al., 2005; O’Keefe and Conway, 1978), odors (Wood et al., 1999), item-place associations (Komorowski et al., 2009), or sounds (Aronov et al., 2017). In future work, we will investigate whether remapping in DQN agents also occurs in scenarios that mimic these more diverse setups.

### 4.2 Spatial and contextual representations in the model network

In our model network, spatially tuned neurons were present more prominently in deeper layers of the neural network, whereas higher layers were more strongly modulated by visual features, suggesting a hierarchical organization of neural codes. Signs of global remapping were found in all layers of the network. While replay had a strong effect on remapping and CR renewal, we found no significant difference in absolute number of spatially tuned cell types.

An interesting difference between deep and superficial layers arises during extinction learning. As mentioned above, spatial representations change during extinction learning and replay helps to maintain the representation of the acquisition context during extinction in context B (see discussion of the results in Fig. 5B). Since in the third experimental phase, trials are not reinforced to test for renewal in context A, the CR is eventually extinguished. To support this change in the behavior in context A, the neural representations have to change, too. Intriguingly, the change does not occur in the superficial layers, where A-A” correlations remain high (Fig. 3B), but they do occur in the deeper layers, where A-A” are peaked around 0. This suggest that the sensory layers retain a stable representation of A, whereas the layers closer to the output have to change their representation to support a different behavior.

In our model, the occurrence of global remapping in the network is strongly correlated with renewal, nevertheless, for low values of the context salience, renewal occurs in the model even though our analysis of remapping does not indicate distinct representations of the two contexts (Fig. 4B). We believe this is an artifact of using the correlations between the populations vectors in the two contexts (A-A’ and A-B) as an indicator of distinct representations. A similar phenomenon has been observed in other studies of memory storage and retrieval, where the similarity between patterns, as measured by a high correlation, did not sufficiently predict whether they would be confused in memory (Neher et al., 2015; Bayati et al., 2018).

While here we used a simplified feed-forward neural network to learn representations, a more detailed model that incorporates high-resolution temporal dynamics will be necessary to account for sub-second neural dynamics and complex neuron responses in hippocampus in the future, e.g. the fast remapping dynamics on the temporal resolution of theta cycles (Jezek et al., 2011), and experience replay (Gillespie et al., 2021).

### 4.3 Testable experimental predictions

The proposed model makes a number of testable experimental predictions. First, our results suggest a close correspondence between remapping and renewal. This prediction could be tested in experiments where neural activity and behavior are recorded simultaneously. Second, we found that the salience of contextual cues plays an important role in extinction and renewal. It has already been shown in experiments that the property of remapping depends on the specificity of the change in experimental conditions Latuske et al. (2018); Leutgeb et al. (2005). It would be interesting to re-analyze these findings in the context of extinction and renewal. Third, we found an important role for replay in the expression of renewal. This finding suggests that disruption of replay Gridchyn et al. (2020) should reliably attenuate renewal.

### 4.4 Conclusion

In summary, we have demonstrated the emergence of context-dependent spatial codes. These emerging representations in our model are compatible with the integrated model and with neural codes in the hippocampus. The occurrence of remapping is strongly correlated with renewal. This suggests that integrated codes are the mechanism for context-dependent extinction and renewal in the DQN agent.

## Acknowledgments

This work was supported by the Deutsche Forschungsgemeinschaft (DFG, German Research Foundation), project number 316803389 (SFB 1280, A14).

